# Identification of biological processes and signaling pathways in lactate-treated cancer cells

**DOI:** 10.1101/2022.02.16.480756

**Authors:** Zhiwen Qian, Hanming Gu, Tingxiang Chang

**Author notes:** Corresponding author: Hanming Gu, SHU-UTS SILC School, Shanghai University, Shanghai, China.

## Abstract

Cancer is a complex disease that involves the alterations of metabolic pathways and tumor microenvironment. Lactate in the tumor microenvironment leads to cancer proliferation, metastasis, and angiogenesis. However, the effect of lactate on prostate cancer cells is still unclear. Here, our objective is to identify the significant molecules and biological processes by analyzing the RNA-seq data. The GSE195639 was produced by the Illumina NextSeq 500 (Homo sapiens). The KEGG and GO analyses show that Herpes simplex virus 1 infection and Rap1 signaling pathway are considered major pathways during the lactate-treated cancer cells. Furthermore, we identified the top ten essential molecules including IL6, CASP3, JUN, MAPK3, BRCA1, PIK3R1, CCNA2, TPI1, APOE, and EXO1. Therefore, our study may provide novel insights into the mechanism of prostate cancers.

## Introduction

Prostate cancer is hard to cure due to the complexity of molecular and cellular heterogeneity^1^. In the USA, high-risk disease contains 15% of all prostate cancer diagnoses^2^. Despite a decreasing death rate over the past 10 years, prostate cancer (PCa) remains the most dominant male malignancy worldwide^3^. Therefore, to meet this clinical need and provide effective treatment methods, it is of utmost importance to study the impact of the tumor microenvironment (TME)^4^.

Lactate was recently identified as a fuel by cancer cells under complete aerobic conditions^5^. Surprisingly, it was shown that lactate resides inside the mitochondria and synthesizes lipids by cancer cells^6^. Moreover, LDHB is localized in the inner mitochondrial membrane and was related to the mitochondrial functions^7^. Thus, lactate is considered to be oxidized to pyruvate in the mitochondria by LDHB^8^. Lactate itself cannot transfer the plasma membrane, so it requires a unique transport mechanism provided by proton-like MCTs^5^. It was also found that lactate shuttle is promoted by pH gradient or by the cellular redox state^9^. The tumor microenvironment includes cancer cells, endothelial cells, cancer-associated fibroblasts, and immune cells^10^. During tumor initiation, neoplastic cells recruit cancer-associated fibroblasts to surround the area through the oxidative stress^11^.

In this study, we identified the signaling pathways of lactate-treated cancer cells by using the RNA sequence data. There were several DEGs, signaling pathways, and protein-protein interaction (PPI) networks identified. The DEGs and PPI networks may guide the treatment of prostate cancer by targeting the tumor microenvironment.

## Methods

### Data resources

Gene dataset GSE195639 was collected from the GEO database. The data was produced by the Illumina NextSeq 500 (Homo sapiens) (The Institute of Oncology Research, Via Vela 6, Bellinzona, Switzerland). The analyzed dataset includes muscles treated by 4 groups of control prostate cancer cells and 4 groups of lactate-treated prostate cancer cells.

### Data acquisition and processing

The data were conducted by the R package as previously described^12–17^. We used a classical t-test to identify DEGs with P<0.05 and fold change ≥1.5 as being statistically significant.

The Kyoto Encyclopedia of Genes and Genomes (KEGG) and Gene Ontology (GO) KEGG and GO analyses were performed by the R package (clusterProfiler) and Reactome. P<0.05 was considered statistically significant.

### Protein-protein interaction (PPI) networks

The Molecular Complex Detection (MCODE) was used to construct the PPI networks. The significant modules were produced from constructed PPI networks and String networks. The signaling pathway analyses were performed by using Reactome (https://reactome.org/), and P<0.05 was considered significant.

## Results

### Identification of DEGs between the prostate cancer cells and lactate-treated prostate cancer cells

To study the impact of lactate on prostate cancer cells, we analyzed the RNA-seq data from the prostate cancer cells treated by lactate. A total of 1351 genes were identified with the threshold of P < 0.01. The top up- and down-regulated genes in the lactate-treated prostate cancer cells were identified by the heatmap and volcano plot (Figure 1). The top ten DEGs were listed in Table 1.

**Figure 1.**
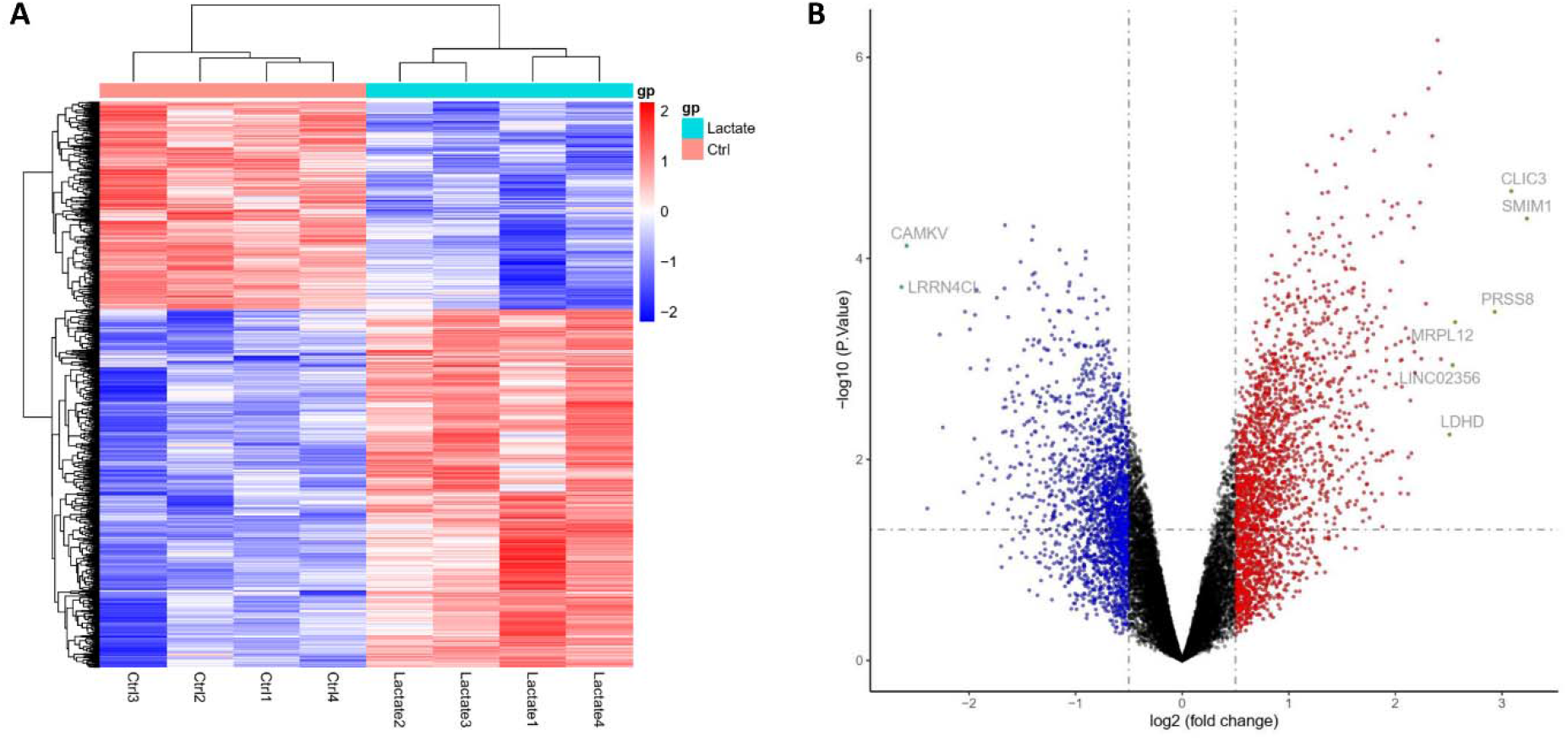
Heatmap and volcano plot were created between the prostate cancer cells and lactate-treated prostate cancer cells. (A) Significant DEGs (P < 0.01) were used to produce the heatmap. (B) Volcano plot for DEGs between the prostate cancer cells and lactate-treated prostate cancer cells. The most significantly changed genes are highlighted by grey dots.

**Table 1.**

| **Entrez gene** | **Gene symbol** | **Fold-change** | **Regulation** |
| --- | --- | --- | --- |
| Top 10 down-regulated DEGs |  |  |  |
| 221091 | LRRN4CL | -2.633055773 | Down |
| 79012 | CAMKV | -2.585151569 | Down |
| 140862 | ISM1 | -2.274053106 | Down |
| 1270 | CNTF | -2.245772454 | Down |
| 10888 | GPR83 | -2.037045662 | Down |
| 10936 | GPR75 | -1.99194638 | Down |
| 79844 | ZDHHC11 | -1.986611188 | Down |
| 170559 | XPOTP1 | -1.948959541 | Down |
| 84519 | ACRBP | -1.943106017 | Down |
| 3798 | KIF5A | -1.928236896 | Down |
| Top 10 up-regulated DEGs |  |  |  |
| 388588 | SMIM1 | 3.233398414 | up |
| 9022 | CLIC3 | 3.087748155 | up |
| 5652 | PRSS8 | 2.932441918 | up |
| 6182 | MRPL12 | 2.560255659 | up |
| 105369984 | LINC02356 | 2.536982989 | up |
| 197257 | LDHD | 2.509296957 | up |
| 136259 | KLF14 | 2.427548018 | up |
| 6676 | SPAG4 | 2.417188052 | up |
| 5745 | PTH1R | 2.396413295 | up |
| 100506211 | MIR210HG | 2.345939024 | up |

### Enrichment analysis of DEGs between the prostate cancer cells and lactate-treated prostate cancer cells

To further determine the potential molecular mechanisms, we performed the KEGG and GO analyses (Figure 2). We identified the top ten KEGG signaling pathways, including “Herpes simplex virus 1 infection”, “Rap1 signaling pathway”, “Biosynthesis of amino acids”, “VEGF signaling pathway”, “Fructose and mannose metabolism”, “Amino sugar and nucleotide sugar metabolism”, “Galactose metabolism”, “Biosynthesis of nucleotide sugars”, “Glycosaminoglycan biosynthesis − chondroitin sulfate / dermatan sulfate”, and “Terpenoid backbone biosynthesis”. We also identified the top ten biological progresses of GO, including “nucleoside phosphate metabolic process”, “hexose metabolic process”, “monosaccharide metabolic process”, “maintenance of location in cell”, “nucleoside diphosphate metabolic process”, “nucleotide phosphorylation”, “maintenance of protein location in cell”, “release of cytochrome c from mitochondria”, “hexose catabolic process”, and “positive regulation of endothelial cell chemotaxis”. We identified the top ten cellular components, including “mitochondrial matrix”, “intrinsic component of organelle membrane”, “integral component of organelle membrane”, “endoplasmic reticulum lumen”, “lysosomal lumen”, “lumenal side of membrane”, “MHC protein complex”, “integral component of lumenal side of endoplasmic reticulum membrane”, “lumenal side of endoplasmic reticulum membrane”, and “synaptic cleft”. We also identified the top ten molecular functions of GO, including “isomerase activity”, “dioxygenase activity”, “pentosyltransferase activity”, “ligand−gated calcium channel activity”, “carbohydrate kinase activity”, “calcium−release channel activity”, “UDP−xylosyltransferase activity”, “xylosyltransferase activity”, “DNA−(apurinic or apyrimidinic site) endonuclease activity”, and “peptidyl−proline dioxygenase activity”.

**Figure 2.**
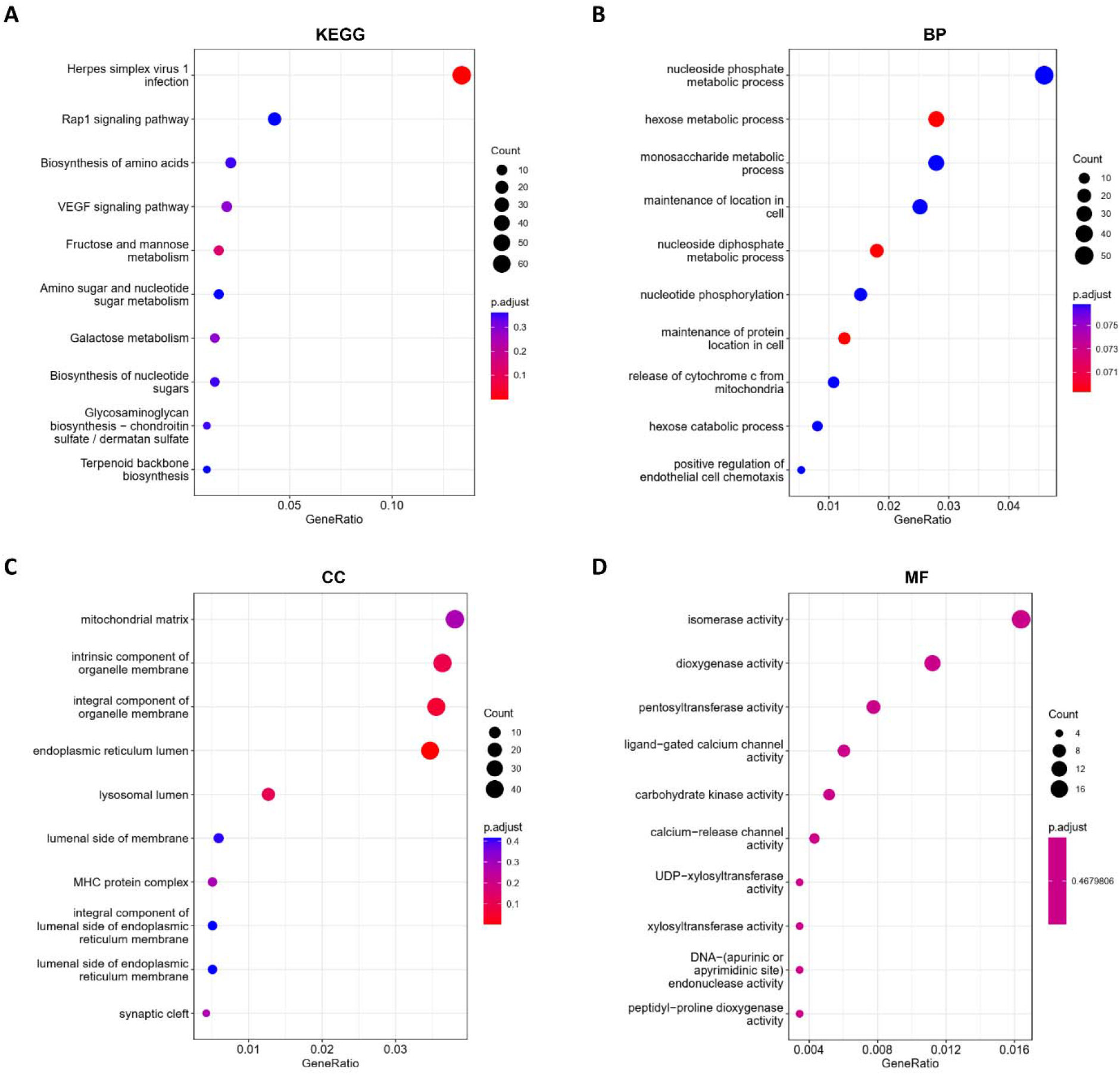
KEGG and GO analyses of DEGs between the prostate cancer cells and lactate-treated prostate cancer cells. (A) KEGG analysis, (B) Biological processes, (C) Cellular components, (D) Molecular functions.

### Construction of PPI network based on the identified DEGs

To understand the potential relationships between the identified DEGs, we created the PPI network by the Cytoscope software. The combined score > 0.2 was set as a cutoff by linking the 108 nodes and 93 edges. Table 2 demonstrated the top ten genes with the highest degree scores. The top two clusters were shown in Figure 3. We further evaluated the DEGs and PPI by Reactome map (Figure 4) and figured out the top ten significant biological processes including “Antigen Presentation: Folding, assembly and peptide loading of class I MHC”, “Endosomal/Vacuolar pathway”, “ER-Phagosome pathway”, “Antigen processing-Cross presentation”, “Interferon gamma signaling”, “Interferon alpha/beta signaling”, “Interferon Signaling”, “Immunoregulatory interactions between a Lymphoid and a non-Lymphoid cell”, “Class I MHC mediated antigen processing & presentation”, and “PD-1 signaling” (Supplemental Table S1).

**Table 2.** Top ten genes demonstrated by connectivity degree in the PPI network.

| **Gene symbol** | **Gene title** | **Degree** |
| --- | --- | --- |
| IL6 | Interleukin 6 | 60 |
| CASP3 | Caspase 3 | 52 |
| JUN | Jun Proto-Oncogene, AP-1 Transcription Factor Subunit | 51 |
| MAPK3 | Mitogen-Activated Protein Kinase 3 | 51 |
| BRCA1 | BRCA1 DNA Repair Associated | 50 |
| PIK3R1 | Phosphoinositide-3-Kinase Regulatory Subunit 1 | 37 |
| CCNA2 | Cyclin A2 | 37 |
| TPI1 | Triosephosphate Isomerase 11 | 36 |
| APOE | Apolipoprotein E | 35 |
| EXO1 | Exonuclease 1 | 34 |

**Figure 3.**
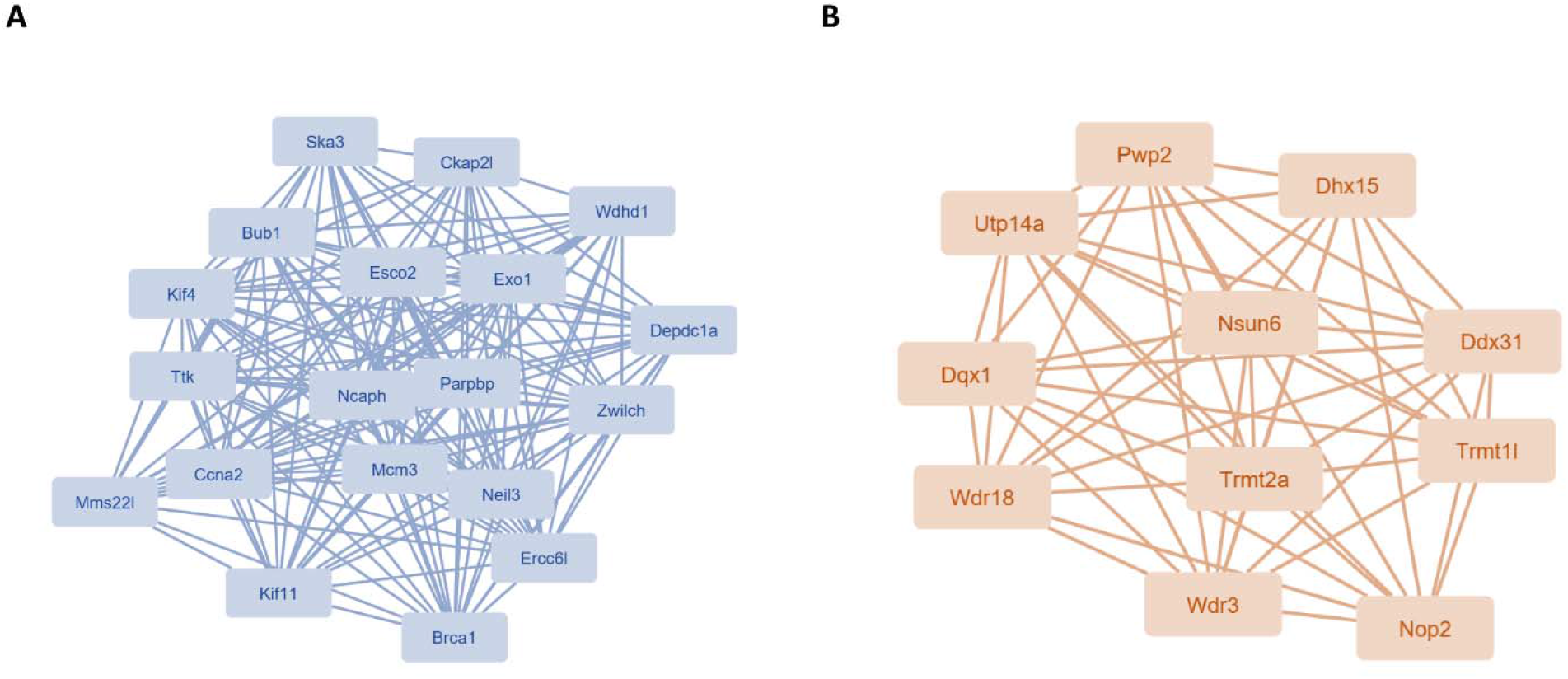
The PPI network analyses of DEGs between the prostate cancer cells and lactate-treated prostate cancer cells. The cluster (A) and cluster (B) were constructed by MCODE.

**Figure 4.**
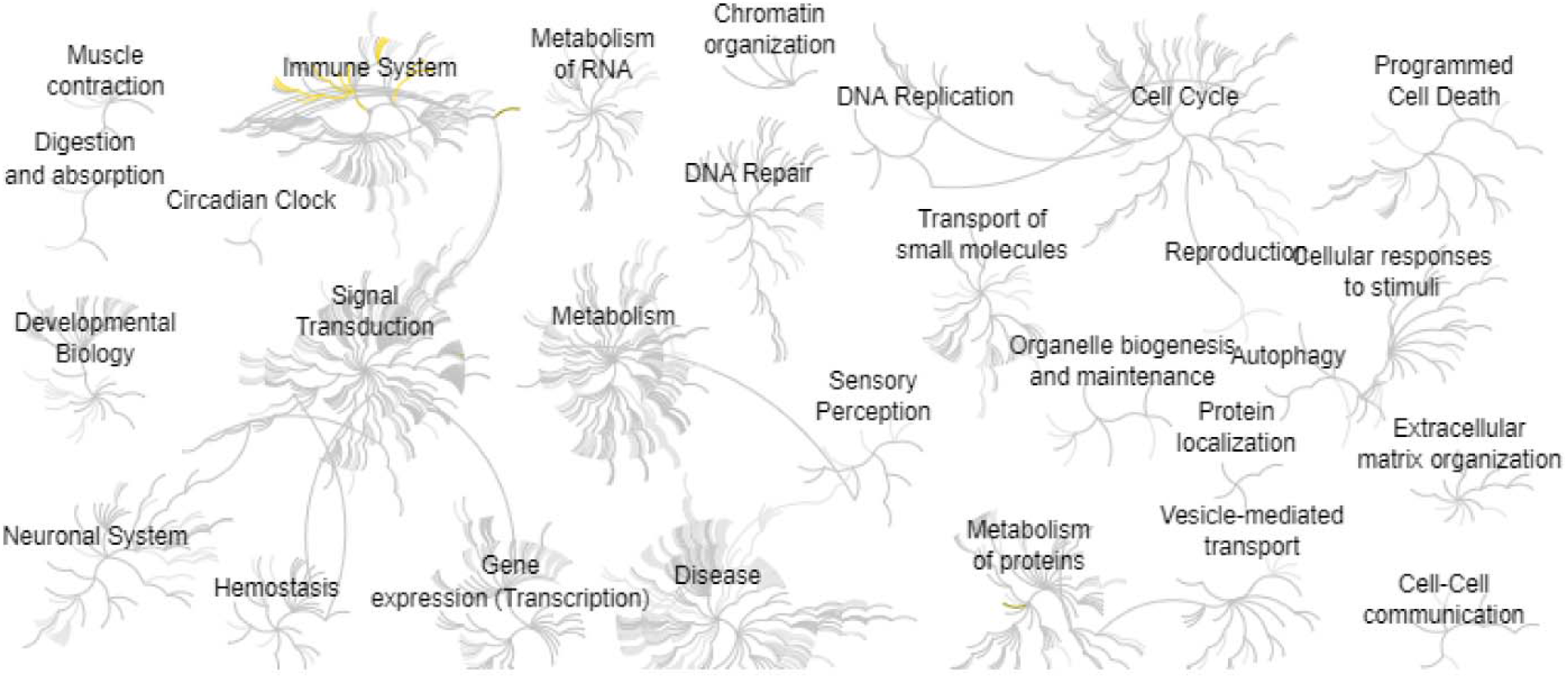
Reactome map representation of the significant biological processes between the prostate cancer cells and lactate-treated prostate cancer cells.

## Discussion

Tumor cells can convert pyruvate into lactate even under normal conditions^18^. Metabolic alterations of tumor cells change the metabolic fluxes, Krebs cycle, and glycolysis, which results in the production of high amounts of lactic acid^19^. The tumor cells can have more lactic acid than normal cells^20^. Thus, understanding the effect of lactate on cancer cells is key to the development of effective therapies.

In this study, Herpes simplex virus 1 infection and Rap1 signaling pathway were considered major pathways during the lactate-treated cancer cells. Oncolytic viruses such as herpes simplex virus type 1 (HSV-1) replicate in tumors and increase immunogenic cell death and induction of immunity^21^. The HSV-1 infection was shown most effective in combination with other anti-cancer agents such as PD1/L1 related immunity^22^. Our study showed that prostate cancer has the same phenotype as HSV-1 infection, suggesting that lactate may also activate cell death or immunity to limit cancer cells^23^. Numerous signaling pathways regulate the activities of G-protein-coupled receptors (GPCRs)^24–26^. GPCRs and Regulators of G protein signaling (RGS) proteins form a feedback loop to mediate the downstream molecules and signaling pathways^27–36^. Among them, GTPase plays different roles in proliferative responses by targeting GPCRs^37^. Rap1 has a similar sequence to Ras, which regulates cell adhesion and integrin function in a variety of cells^38^. Specifically, Rap1 initiates ERK signaling that is activated in many malignancies and is an effective target of cancers^39^.

In addition, we also discovered the top ten interaction molecules by constructing the PPI network. Takuji Hayashi et al. found that the high-fat diet can induce inflammation and promote prostate cancer growth through IL6/pSTAT3 signaling in the mice^40^. Circadian clocks regulate a variety of biological processes including inflammation, metabolism and aging^41–51^. I Aiello et al. found that the disruption of circadian clock can affect the IL6 expression and change the tumor state^52^. Shu-Pin Huang and Chun-Hsiung Huang et al. found that the genetic variants in CASP3 may affect the survival in prostate cancer patients receiving androgen-deprivation therapy by analyzing several genome-wide association studies (GWAS)^53^. Christine Moore demonstrated that the upregulation of JUN is critical to the differential effect on apoptosis in prostate cancer cells^54^. Rong Deng et al. found that MAPK1 mediates the ULK1 degradation to decrease mitophagy and promotes cancer cell metastasis^55^. Genomic analysis indicates BRC1 is strongly associated with prostate cancer risk and aggressiveness^56^. PI3K regulatory subunit gene PI3K1 is inhibited in prostate cancers^57^. Weighted gene co-expression network indicates CCNA2 is a potential therapeutic target for prostate cancer by inhibiting cell cycle^58^. TPI1 is associated with the tumor microenvironment and squamous cell cancer^59^. APOE polymorphism affects aggressive behavior in prostate cancer cells by inhibiting cholesterol homeostasis^60^. Fei Luo and Kuo Yang et al. found the expression of EXO1 is related to clinical progression, metastasis, and survival prognosis of prostate cancer^61^.

In conclusion, this study discovered the biological functions of lactate in prostate cancer cells. Herpes simplex virus 1 infection signaling and Rap1 signaling pathway are the major pathways during the lactate-treated cancer cells. Our study may help to shed light on the mechanism of tumor metabolism.

## Author Contributions

Zhiwen Qian, Tingxiang Chang: Methodology. Hanming Gu: Conceptualization, Writing and Editing.

